# Phycosphere-associated bacteria differentially impact accessibility of dust-bound iron to model diatom *Phaeodactylum tricornutum*

**DOI:** 10.64898/2026.06.30.735391

**Authors:** Nicole R. Coffey, Brooke N. Newell, Kieran Manning, Kristina A. Rolison, Xavier Mayali, Rhona K. Stuart, Rene M. Boiteau

## Abstract

In marine ecosystems, phytoplankton growth is frequently limited by iron, a micronutrient, due to its poor solubility from major sources such as atmospheric dust. Many phytoplankton cannot access dust-bound iron independently, and processes that solubilize this iron remain poorly understood. Here, we investigated whether bacterial partners can enhance phytoplankton growth under iron-limited conditions by facilitating utilization of dust-bound iron. Our study focused on *Phaeodactylum tricornutum*, a model diatom that is adapted to low iron growth conditions, grown in co-culture with bacteria isolated from its phycosphere. In iron-limited experiments using mineral dust as the sole iron source, the addition of *Marinobacter* significantly enhanced diatom growth compared to axenic controls, whereas *Stappia* significantly suppressed it. However, under iron-replete conditions, neither bacterium affected growth. These results indicated that under low-iron conditions, *Marinobacter* actively alleviates iron deficiency. Co-cultured bacterial cell abundances remained at least an order of magnitude lower than diatom cells. *Marinobacter* also enhanced algal growth within days of dust addition to established Fe-limited co-cultures, indicating its beneficial effect on *P. tricornutum* was not unique to a system in which it was newly introduced. Exometabolomic profiling comparing the axenic diatom and co-cultures revealed a suite of condensed aromatic organosulfur and peptide-like compounds associated with bacterial presence, as well as compounds that appeared to be unique to each co-culture, hinting at a molecular underpinning of each strain’s impact. Our findings demonstrate that low-abundance members of the phycosphere community can have a significant impact on host growth by modulating the accessibility of dust-bound Fe.

## Introduction

Iron (Fe) is an essential micronutrient in marine ecosystems [1], but is present in extremely low concentrations through much of the surface ocean [2,3]. As a result, 30-40% of surface ocean primary production is limited by its availability [4,5]. The prominent source of Fe to surface waters over much of the ocean is aeolian dust [6–8], though this dust delivery is ephemeral in space and time [6,9]. Furthermore, dust-derived Fe has low solubility in seawater [10,11]. This poses a challenge to many marine microorganisms, particularly phytoplankton, that depend on aeolian Fe but lack strategies to solubilize Fe from mineral dust themselves. Although mutualistic relationships between major marine phytoplankton that fix carbon and associated heterotrophic taxa that can solubilize Fe have been speculated [12], there is currently little evidence that phycosphere heterotrophs can improve diatom access to dust-bound Fe or significantly increase primary production.

Biological strategies for solubilizing Fe from mineral dust in seawater follow one of two mechanisms: siderophore production; or reductive dissolution, in which mineral-bound Fe(III) is reduced to more soluble but less stable Fe(II). Siderophores are small biomolecules produced in response to Fe limitation to enhance Fe bioavailability and uptake. Though some eukaryotic algae have transporters that enable them to utilize exogenous siderophores for Fe uptake, they are unable to produce these molecules themselves [12,13], implying a dependence on bacterial members of the community for Fe acquisition. In culture and incubation experiments, diatoms and haptophytes from chronically Fe-limited waters have been shown to take up Fe bound to bacterially-produced siderophores such as desferrioxamine B [14,15]. Photodegradation of siderophore-Fe complexes, including those produced by strains of *Marinobacter*, has also been demonstrated to enhance algal Fe uptake [16,17]. Bacteria associated with diatom and haptophyte blooms in these regions have been shown to have the genetic capacity for siderophore biosynthesis [18,19], suggesting that these compounds may play a role in regulating Fe bioavailability. Though the cyanobacterium *Trichodesmium* and its associated bacteria have been shown to utilize siderophores [20] and are suspected to mobilize mineral-bound Fe via reductive dissolution [21–23] to enhance *Trichodesmium*’s Fe uptake, the mechanisms of how diatom-bacteria interactions may enable (or inhibit) access to mineral-bound Fe are currently unclear.

Diatom cellular response to Fe limitation has been extensively characterized, providing a useful model to investigate diatom-bacteria interactions on particulate Fe acquisition. In particular, *Phaeodactylum tricornutum*, a coastal diatom [24–26], shares transcriptional responses to Fe limitation with several open ocean species [27,28], making it a useful model for studying Fe deficiency in phytoplankton [29–33]. Previous culture experiments have shown *P. tricornutum* growth and photosynthetic efficiency are enhanced with Fe(II)-rich primary silicate mineral additions [34]. However, Fe(III)-rich oxyhydroxide minerals are more difficult for the diatom to access due to poorer solubility [34], and may promote reliance on members of the bacterial community for Fe acquisition. As Fe-oxide minerals are a significant Fe-bearing component of natural dust sources [34–36], understanding reactions to mineral addition is crucial for prediction of system-level adaptation to changing Fe inputs. *P. tricornutum* harbors putative siderophore receptors and has been shown to take up exogenous hydroxamate siderophores, but is unable to produce these Fe acquisition molecules [12,37]. Further, *P. tricornutum* is known to influence its surrounding bacterial community under Fe replete conditions through a variety of characterized mechanisms, such as specific exometabolite excretion [38–40].

In this study, we set out to characterize how *P. tricornutum*-bacteria interactions respond to Fe limitation to ultimately better understand the role of these interactions in regulating particulate Fe acquisition and its allocation between different community members. First, we used a co-culture of *P. tricornutum* with its associated bacterium *Marinobacter* sp. (strain 3-2) or *Stappia* sp. (strain ARW1T) and a liquid chromatography electrospray ionization mass spectrometry approach to profile the exometabolome of axenic diatom cultures and co-cultures under high and low dust treatments. *Marinobacter* was isolated from *P. tricornutum*’s phycosphere [40,41], and has previously been shown to positively impact the diatom’s growth, significantly increasing algal cell density and lipid content relative to axenic controls [41]. It is also suspected to produce a siderophore-like molecule or redox active metabolites based on antiSMASH [42] genome annotations (Fig. S1), suggesting it is a promising candidate for a diatom beneficial partner. Though *Stappia* was also isolated from *P. tricornutum*’s phycosphere [38,40] and is suspected to produce a siderophore-type molecule (Fig. S1), it was not demonstrated to increase algal biomass in Fe-replete co-culture conditions [40], potentially providing an example of a commensal partner. To evaluate the effect of *Marinobacter* on dust dissolution in habituated co-culture with *P. tricornutum*, we then performed a second experiment in which dust was added to low Fe-acclimated cultures during early exponential growth phase to mimic a dust deposition event. Our results demonstrate that 1) the presence of *Marinobacter*, even in low abundance, significantly impacted the diatom’s growth under the low dust treatment and after dust deposition events; 2) *P. tricornutum*’s exometabolome significantly shifted in response to both beneficial and antagonistic bacteria; and 3) *Marinobacter* fostered the accumulation of four major suites of compounds that may play a role in Fe solubilization and/or algal growth. These results provide novel insight into *P. tricornutum*’s interaction with two phycosphere-associated bacteria and suggest that certain suites of non-siderophore compounds may be important for regulating community Fe acquisition.

## Materials and Methods

### Culture Conditions and Sample Generation – Dust as the Sole Fe Source

Axenic stock cultures of *P. tricornutum* (Bohlin strain CCMP 2561) in modified f/2 medium [43,44], with 1 µM Fe-EDTA, which has previously been demonstrated to be an iron replete condition for this diatom [45] (Fig. S2). These cultures were incubated at 19.0°C under cool white fluorescent lighting (2030 lx) on a 12-hour light/dark cycle in an algal growth chamber (Percival Scientific). Axenic algal cultures were checked for bacterial contamination at each transfer by streaking on marine broth 2216 (BD Difco) agar plates and monitoring for bacterial growth.

To pre-treat the diatom for experimental conditions, axenic *P. tricornutum* was transferred to fresh medium with Fe added as 0.018 mg Arizona Test Dust (AZTD [46]), targeting a dissolved Fe concentration of 10 nM based on solubility estimates derived from previous acid mobilization studies [35]. This concentration reflects previous work establishing 5-10 nM Fe as limiting for this diatom [29,45,47], and represents the “low dust” treatment in this study. All AZTD used for culture medium was sterilized in a 100% ethanol suspension for 30 minutes, then rinsed with sterile water and suspended in sterile medium before use. The pre-treated parent cultures were allowed to grow until they reached a cell density of ∼10^5^ cells/mL, and were then transferred 1:100 to fresh low dust medium.

To prepare both *Marinobacter* and *Stappia* for experimental conditions, a single colony was picked from marine broth 2216 (BD Difco) agar plates and grown in 5 mL liquid marine broth. To minimize Fe carryover from the bacterial stock, the culture was centrifuged at 1440 rcf for 5 minutes. The supernatant was removed, and the pellet was rinsed with 3 mL sterile f/2 medium with no Fe added. This suspension was centrifuged at 3500 rpm for 5 minutes, and the supernatant was discarded. The rinsed pellet was then resuspended in 5 mL sterile f/2 medium for inoculation into algal flasks.

After two passages through the low dust pre-treatment, the diatoms were transferred to the experimental conditions. 100 mL FeCl_3_-free f/2 medium in 250 mL polycarbonate flasks were inoculated to a starting cell density of 10^3^ algal cells/mL from the pretreated parent culture for both the low (0.018 mg) and high (1.8 mg, target soluble Fe concentration of 1 μM) dust treatments. Axenic *P. tricornutum* cultures received no further amendments. For co-cultures with bacterial strains, 100 μL of the f/2 bacterial pellet suspension were added to the treatment flasks, targeting a starting cell density of 10^4^ bacterial cells/mL (actual load: 4.1 × 10^3^ *Marinobacter* cells/mL and 1.1 × 10^2^ *Stappia* cells/mL). All treatments were prepared in triplicate. Cultures were monitored by algal cell counts using a hemocytometer. Samples were periodically fixed for bacterial cell counts by removing 1.5 mL subsamples of each culture and adding 30 μL 25% glutaraldehyde and incubating for 10-minutes in the dark before flash freezing in liquid nitrogen and stored (−80 °C) until analysis. Bacterial cell counts were conducted by flow cytometry (GUAVA EasyCyte Plus flow cytometer and Guava CytoSoft (v5.3), Millipore) after staining with SYBR Green I (Life Technologies) following previously established methods [48,49]. Subsamples (10 mL) were collected and filtered (0.22 μm, Whatman) for exometabolomic analysis after 14 (t_14_) days of growth.

### Culture Conditions and Sample Generation – Dust Addition Experiments

Axenic diatom stock cultures were maintained as above. To prepare an established *Marinobacter*-*P. tricornutum* co-culture for these experiments, parent *P. tricornutum* cultures were inoculated with *Marinobacter* as above, and passaged eight times in the modified f/2 media before beginning pre-treatment for the low Fe experiment. To pre-treat both axenic *P. tricornutum* and *Marinobacter* co-cultures for the experiment, both cultures were transferred to fresh medium with 10 nM Fe added to an algal cell density of 5000 cells/mL. The pre-treated parent cultures were allowed to grow until they reached a cell density of ∼10^5^ cells/mL, and were then transferred to fresh low dust medium at a final cell density of 5000 algal cells/mL.

After two passages through the low Fe pre-treatment, the cultures were transferred to the experimental conditions. 100 mL f/2 medium (10 nM Fe) in 250 mL polycarbonate flasks were inoculated to a starting cell density of 5000 algal cells/mL from the pretreated parent cultures in triplicate. Cultures were maintained under the same incubation conditions as the parents and monitored by fluorescence. All cultures were spiked with 0.018 mg AZTD (∼10 nM soluble Fe addition). Immediately before and after dust addition, Fe(II) was measured using a chemiluminescent method (details in Supplementary Methods).

### Exometabolite Extraction

Ten mL filtered subsamples from the experiment with dust as the only Fe source were solid phase extracted without pH adjustment using 0.1 g PPL cartridges (Agilent Technologies) using previously established methods [45,50,51]. Briefly, these cartridges were prepared using sequential 3 mL flushes of LC-MS grade methanol (VWR Chemical) 0.1% trace metal-grade HCl (Fisher Chemical), and ultrapure water prior to sample loading. After sample loading, cartridges were rinsed with another 3 mL ultrapure water to remove residual salts and frozen (−20°C) until elution. To prepare for analysis, thawed columns were eluted with 1.5 mL LC-MS grade methanol and dried down to approximately 50 μL. This concentrate was then diluted with ultrapure water to a final volume of 1 mL. A 500 μL aliquot of each sample was transferred to a 2 mL polypropylene HPLC vial (VWR) and spiked with an internal standard (1 μM cyanocobalamin, Sigma Aldrich). Pooled samples for quality control were generated by combining an equal volume of each sample.

### Liquid Chromatography Electrospray Ionization Mass Spectrometry (LC-ESI-MS)

Samples were separated by high pressure liquid chromatography (Ultimate 300 RSLCnano, Thermo Scientific). Full loop (50 μL) injections of each sample were loaded onto a C18 column (0.5 × 100 mm, 3.5 μm particle size; Agilent Zorbax XDB-C18) at 30 μL/ min (solvent A, 95% ultrapure water + 5 mM ammonium formate; solvent B, 5% methanol + 5 mM ammonium formate). Separations were performed holding the column at 40°C over a 30-minute gradient from 95% solvent A and 5% solvent B to 5% solvent A and 95% solvent B, followed by a 5-minute hold at 95% solvent B. Eluent composition was then returned to 95% solvent A and held for 9 minutes prior to introducing the next sample. All chemicals and solvents were LC-MS grade (Optima; Fisher Scientific). Consistency in the peak area of the internal standard cyanocobalamin (4.9% standard deviation) indicated analytical stability over the course of the run.

Flow from the LC was directed into a Thermo Scientific Orbitrap IQ-X mass spectrometer equipped with a heated ESI source set to a capillary voltage of 3500 V, sheath, auxiliary and sweep gas flow rates of 5, 2, and 0 (arbitrary units), and ion transfer tube and vaporizer temperatures of 275 °C and 75 °C. MS^1^ scans were collected over a *m/z* range of 100–1000 in high resolution (500 k at *m/z* 200, transient length 1024 ms) positive mode. MS^2^ fragmentation spectra were collected in the ion trap using data dependent acquisition with an HCD energy of 30, a precursor quadrupole isolation width of 1.60, and at an acquisition rate of 35 spectra per second. Dynamic exclusion with a duration of 5 s and a mass tolerance of 10 ppm was used to limit repeated fragmentation of the same precursor.

### Exometabolome Feature List Generation

A python-based workflow leveraging the open-source platform CoreMS [52] and CoreMS tools [53] was used to generate a molecular formula assigned metabolite feature list using the LC-ESI-MS data set based on previously established methods [54]. For each analysis, mass spectra were averaged over 2-minute time intervals across the chromatographic gradient and internally calibrated using a series of ubiquitous polysiloxane masses to improve mass accuracy. Molecular formulae were assigned to these calibrated masses using search criteria consistent with most biologically relevant molecules [55–57]: C 1-40, H 4-80, O 0-16, N 0-8, S 0-2, with a maximum double bond equivalents (DBE) of 20. If multiple assignments could be made to a peak within the mass error tolerance, the assignment with the highest confidence score was selected. To generate the feature list, all assigned features in each sample were aligned using feature *m/z* and retention time interval to allow for intensity comparisons across the dataset, resulting in a final list of 9,465 features with confidently assigned molecular formula after blank subtraction.

### Metabolite Structural Annotation

Structural annotation of metabolites were obtained using tandem MS-MS fragmentation spectra library matching in MSDial version 5.5.24113 [58,59] and GNPS2 release 2025.02.12 [60,61]. Only features with intensities above 1,000 were considered. Mass tolerances were constrained by 0.01 Da and 0.1 Da for MS^1^ and MS^2^ spectra, respectively. An identification score cutoff of 70% was imposed to limit false matches to the Pos VS17 authentic standard spectral library in MSDial (324,191 entries). Additional structural annotations were obtained by feature based molecular networking workflow in GNPS2. Feature processing was conducted using the following parameters: precursor ion tolerance of 5, fragment ion tolerance of 0.2, minimum cosine score of 0.7, minimum of 4 matched peaks, top K of 10, and a maximum component size of 100. For the GNPS2 library search, matches were restricted to a minimum cosine score of 0.5 and a minimum of 4 matched peaks.

## Results

### Bacterial Co-incubation Significantly Impacted Algal Growth Under Low Dust Treatment

We first conducted experiments using a single Fe limited axenic *P. tricornutum* parent culture with dust as the only Fe source, and introduced one of two bacterial strains (*Marinobacter* and *Stappia*) at the start of the incubation with the goal of establishing differences in diatom growth and exometabolite profiles due to the presence of different bacterial partners. By starting with the same parent diatom culture, the specific effects of different bacterial partners could be evaluated. The co-culture with *Marinobacter* led to significantly increased *P. tricornutum* growth rate and yield under Fe limitation relative to the axenic control. Fe limited and replete growth under the low and high dust treatments respectively were consistent with previous published and in-house experiments using Fe-EDTA as an iron source, which reported Fe limitation of *P. tricornutum* growth at dissolved Fe concentrations between 5 and 10 nM [29,45,47] (Fig. S2).

Under the low dust treatment, significant differences in algal cell abundance between axenic *P. tricornutum* and co-cultures with *Marinobacter* emerged after 4 days of growth (non-paired heteroscedastic t-test, p < 0.05), with the algal cell density at the final time point being nearly 6-fold higher in the co-culture than in the axenic diatom culture (Fig. 1A). In the high dust treatment, final algal cell concentrations under the high dust treatment did not differ significantly between biological treatments (Fig. 1B). Comparing high and low dust treatments, the low dust treatment led to Fe limitation with significant differences in cell density emerging after 4 (p < 0.005) and 7 (p < 0.005) days of growth, respectively (Fig. 1A and B). Diatoms in co-culture with *Marinobacter* were still Fe limited under the low dust treatment, but reached 18% of the final cell density of the co-culture under the high dust treatment.

**Figure 1.**
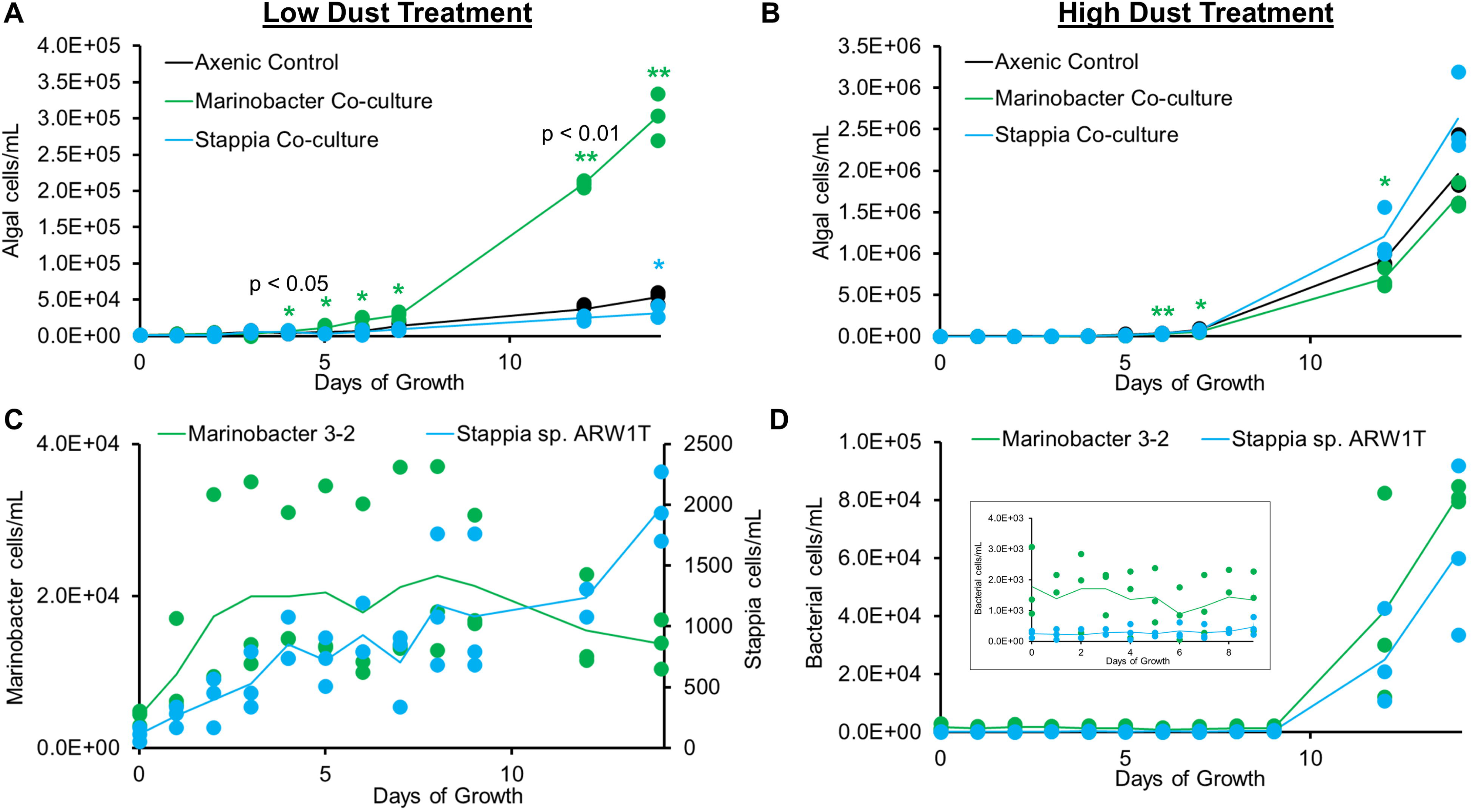
Algal cell abundances (cells/mL) in low dust (A) and high dust (B) treatments of axenic *P. tricornutum* (black) and in co-culture with *Marinobacter* (green) and *Stappia* (blue). Individual replicates are plotted as points, while the line represents the average (n=3). Significant differences between algal cell abundance are noted marked “*” for p < 0.05 and “**” for p < 0.01(unpaired heteroscedastic t-test). Bacterial cell abundances (cells/mL) of *Marinobacter* (green) and *Stappia* (blue) in the low dust treatment (C) and high dust treatment (D) in co-culture with *P. tricornutum*. Inset shows bacterial populations over the first 9 days of growth.

Co-cultivation with *Stappia*, in contrast to *Marinobacter*, inhibited algal growth in the low dust treatment. After 14 days in the low dust treatment, co-culture algal abundance was significantly lower, reaching ∼59% of the axenic *P. tricornutum* control (p < 0.05). *Stappia*’s significant inhibitory effect persisted through 26 days of growth (Fig. S3). Under the high dust treatment, however, there was no significant difference in algal cell abundance between the *Stappia* co-culture and the axenic control, consistent with previous co-culture studies under Fe replete conditions [40].

Despite the significant effect of bacteria on algal growth in Fe limited conditions, bacterial cell abundance remained relatively low throughout the experiment, with *Marinobacter* reaching final cell abundances of 1.4 ± 0.3 ×10^4^ cells/mL and 8.2 ± 0.3 ×10^4^ cells/mL in the low (Fig. 1C) and high (Fig 1D) dust treatment, and *Stappia* reaching a final cell density of 2.0 ± 0.3 × 10^3^ cells/mL and 6.2 ± 3 × 10^4^ cells/mL in the low and high dust treatments, respectively. In the low dust treatment, *Marinobacter* biomass increased modestly early in the experiment before the diatom entered the log phase of growth, and remained relatively constant as the algae grew. *Stappia* bacterial abundance, however, gradually increased over the 14-day experiment under the low dust treatment. Conversely, under the high dust treatment, bacterial cell counts in both co-cultures remained relatively constant until after 9 days of growth, when the diatom was already in the log phase of growth. The bacterial cell density then increased as the diatom cell abundance increased, with no significant difference in bacterial abundance (p < 0.05) throughout the experiment except at 3 days of growth (p = 0.04).

### Enhanced Growth from Dust Addition to Bacterial Co-incubations

Having established that *Marinobacter* enhances *P. tricornutum* growth with dust, we next aimed to investigate Fe mobilization in established co-cultures experiencing dust deposition. After acclimating the *Marinobacter*-diatom coculture to Fe limited conditions, we monitored the growth before and after dust addition. This mimics the natural atmospheric deposition events, where microbial communities under Fe stress upregulate Fe acquisition strategies. The pre-established co-culture with *Marinobacter* led to significantly increased *P. tricornutum* growth under low Fe conditions prior to dust addition relative to the axenic control (p < 0.05), with the co-culture reaching a fluorescence approximately 1.4 times higher than that of the axenic culture after 7 days of growth (Fig. 2). This enhancement was likely attributed to *Marinobacter* increasing the uptake of originally added Fe by *P. tricornutum*. After the dust addition at 7 days of algal growth, the *Marinobacter* co-culture continued to grow significantly, with fluorescence increasing by ∼2-fold relative to day 7 (p < 0.01) after 10 and 11 days of growth (3 and 4 days post-dust addition, respectively) than before dust addition at 7 days of growth. This was not seen in the axenic control, which did not show significantly increased growth after dust addition, based on a comparison of fluorescence at the end of the experiment relative to day 7. These results further demonstrate the beneficial effect of *Marinobacter* under iron-limited conditions and suggest that acclimated co-cultures of *P. tricornutum* with *Marinobacter* are better able to utilize dust-bound iron than the axenic diatom culture.

**Figure 2:**
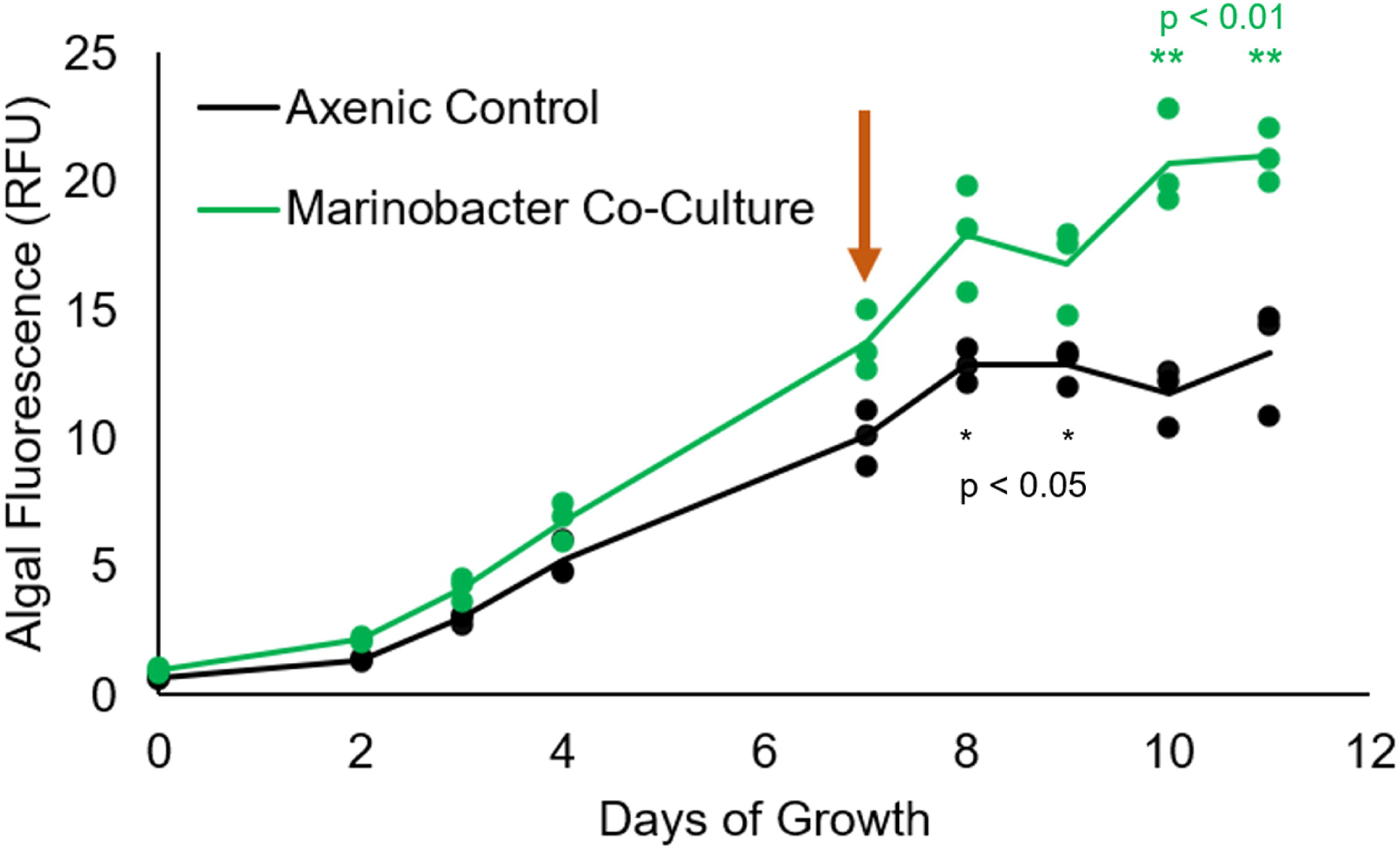
Time series of algal fluorescence (relative fluorescence units) in axenic *P. tricornutum* (black) and co-culture with *Marinobacter* (green). Individual replicates are plotted as points, while the line represents the average of 3 replicates for each treatment. Time of dust addition (7 days) is noted with a rust-colored arrow. Significant differences between algal fluorescence at 7 days of growth and post-dust addition (unpaired heteroscedastic t-test) are marked “*” for p < 0.05 and “**” for p < 0.01.

### Exometabolomic Fingerprint of Bacterial Impacts on Diatom Growth

To investigate exometabolites that potentially contributed to the growth effect of the phycosphere bacteria, we investigated changes in *P. tricornutum*’s exometabolome in response to shifting Fe availability and the presence of bacterial partner *Marinobacter*. LCMS analysis annotated 9,465 distinct exometabolites, revealing major differences between axenic and co-culture growth conditions (Fig. S4).

To further investigate which of these compounds could be contributing to the near-sixfold increase in the diatom’s growth under the low dust treatment with *Marinobacter* or the 41% reduction in algal growth with *Stappia*, we examined a curated list of the 15 metabolites with greatest measured intensity of those significantly (≥2-fold difference, Bonferroni adjusted two-tailed t-test p value < 0.05) enriched in the *Marinobacter* co-culture in the initial CoreMS analysis, as well as an additional 11 curated features that were 1) significantly enriched in the low dust *Stappia* co-culture relative to the axenic control or 2) significantly enriched in the low dust *Marinobacter* co-cultures, but not annotated with molecular formula using CoreMS (Fig. 3, Table 1). These compounds fell into five overarching categories: 1) those associated with higher diatom cell density, as indicated by their presence in the high dust axenic diatom cultures; 2) those associated with higher diatom cell density, but more abundant in the presence of either bacterial strain relative to the axenic control; 3) those with higher intensity in bacterial co-cultures, regardless of dust treatment; 4) features that appear to be unique to *Marinobacter* co-cultures; and 5) a feature that is unique to the *Stappia* co-cultures.

**Figure 3:**
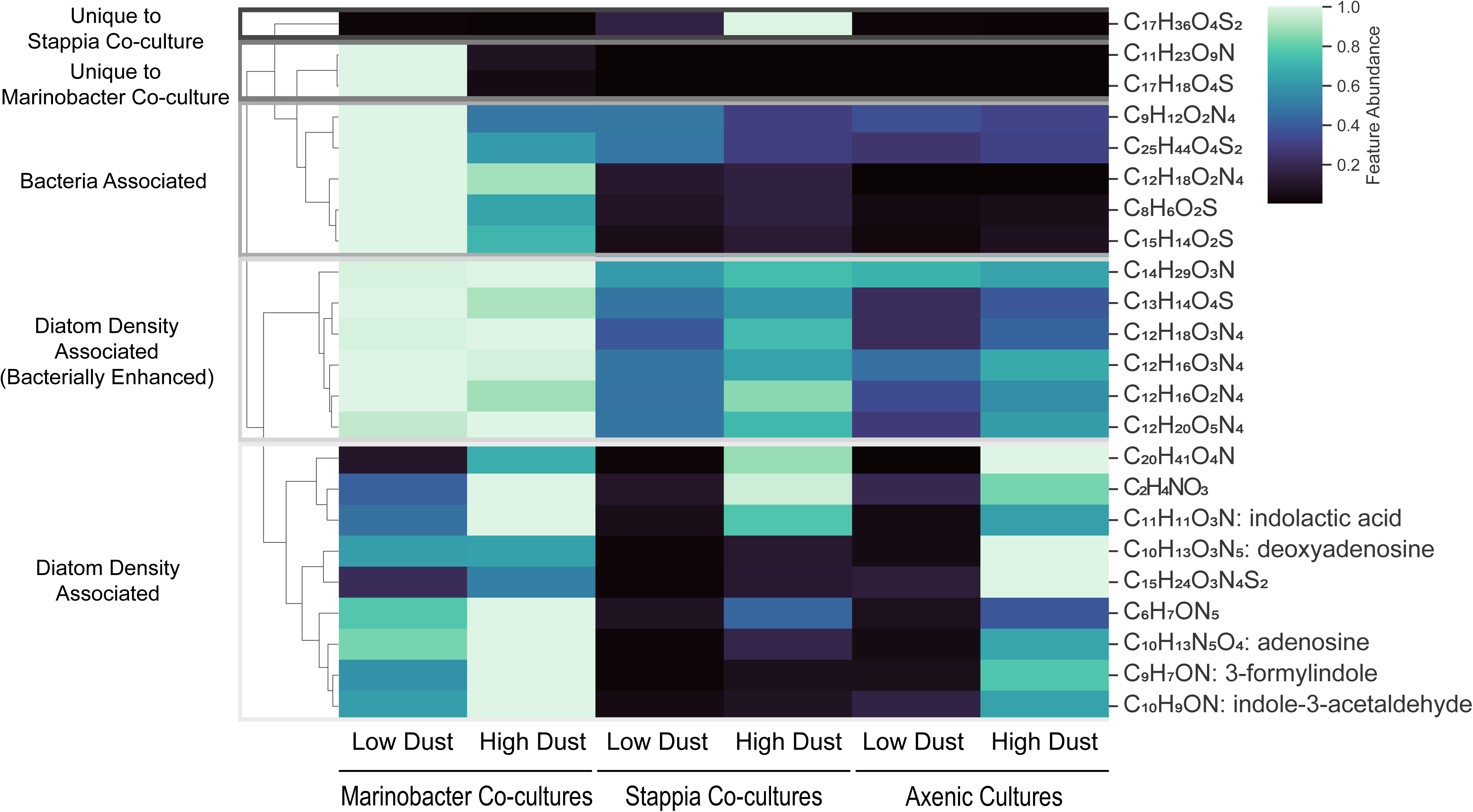
Heat map of the metabolites with the highest relative abundance that were significantly (p-adj < 0.05) higher in low dust co-cultures of *P. tricornutum* with *Marinobacter* relative to axenic cultures or with *Stappia* relative to axenic cultures. Putative structural annotations are indicated.

**Table 1:** Formula assignments and putative structural annotations of the most intense features that were significantly (p-adj < 0.05) higher in low dust cultures with *Marinobacter* relative to axenic cultures or with *Stappia* relative to axenic cultures. *Formula assigned using ChemCalc [62]

| *m/z* | RT<br>(min) | Molecular<br>Formula | *m/z* error<br>(ppm) | MS/MS Annotation | Cosine Similarity<br>(GNPS2) | Match Score<br>(MSDial) |
| --- | --- | --- | --- | --- | --- | --- |
| 113.0084 | 2.90 | *C<sub>2</sub>H<sub>4</sub>O<sub>3</sub>N<br>(Na<sup>+</sup> adduct) | +0.320 | - | - | - |
| 146.0602 | 10.62 | C<sub>9</sub>H<sub>7</sub>O N | +0.035 | 3-formylindole | - | 1.8639 |
| 160.0757 | 10.35 | C<sub>10</sub>H<sub>9</sub>O N | -0.064 | indole-3-acetaldehyde | 0.8786 | - |
| 166.0722 | 3.71 | C<sub>6</sub>H<sub>7</sub>O N<sub>5</sub> | -0.025 | - | - | - |
| 167.0160 | 15.85 | C<sub>8</sub>H<sub>6</sub>O<sub>2</sub>S | +0.057 | - | - | - |
| 206.0812 | 8.34 | C<sub>11</sub>H<sub>11</sub>O<sub>3</sub>N | +0.178 | indolactic acid | - | 1.212 |
| 209.1032 | 4.59 | C<sub>9</sub>H<sub>12</sub>O<sub>2</sub>N<sub>4</sub> | +0.306 | - | - | - |
| 249.1344 | 7.72 | C<sub>12</sub>H<sub>16</sub>O<sub>2</sub>N<sub>4</sub> | +0.245 | - | - | - |
| 251.1501 | 8.22 | C<sub>12</sub>H<sub>18</sub>O<sub>2</sub>N<sub>4</sub> | +0.331 | - | - | - |
| 252.1092 | 5.26 | C<sub>10</sub>H<sub>13</sub>O<sub>3</sub>N<sub>5</sub> | +0.317 | deoxyadenosine | 0.9990 | - |
| 259.0788 | 20.67 | C<sub>15</sub>H<sub>14</sub>O<sub>2</sub>S | +0.317 | - | - | - |
| 260.2219 | 22.20 | C<sub>14</sub>H<sub>29</sub>O<sub>3</sub>N | +0.202 | - | - | - |
| 265.1296 | 5.60 | C<sub>12</sub>H<sub>16</sub>O<sub>3</sub>N<sub>4</sub> | +0.199 | - | - | - |
| 267.0688 | 15.74 | C<sub>13</sub>H<sub>14</sub>O<sub>4</sub>S | +0.395 | - | - | - |
| 267.1449 | 8.80 | C<sub>12</sub> H<sub>18</sub> O<sub>3</sub> N<sub>4</sub> | +0.293 | - | - | - |
| 268.1043 | 5.00 | C<sub>10</sub> H<sub>13</sub> O<sub>4</sub> N<sub>5</sub> | +0.396 | adenosine | 0.9983 | - |
| 301.1506 | 5.85 | C<sub>12</sub> H<sub>20</sub> O<sub>5</sub> N<sub>4</sub> | +0.098 | - | - | - |
| 314.1447 | 3.60 | C<sub>11</sub> H<sub>23</sub> O<sub>9</sub> N | -0.272 | - | - | - |
| 319.1001 | 3.56 | C<sub>17</sub> H<sub>18</sub> O<sub>4</sub> S | +0.015 | - | - | - |
| 360.3110 | 19.98 | C<sub>20</sub> H<sub>41</sub> O<sub>4</sub> N | +0.054 | - | - | - |
| 369.2131 | 8.15 | C<sub>17</sub> H<sub>36</sub> O<sub>4</sub> S<sub>2</sub> | +0.994 | - | - | - |
| 373.1361 | 10.96 | C<sub>15</sub> H<sub>24</sub> O<sub>3</sub> N<sub>4</sub> S<sub>2</sub> | -0.219 | - | - | - |
| 473.2748 | 22.54 | C<sub>25</sub> H<sub>44</sub> O<sub>4</sub> S<sub>2</sub> | -0.231 | - | - | - |

Two features (C_11_H_23_O_9_N, C_17_H_18_O_4_S) were associated with the presence of *Marinobacter* and were not detected in the axenic control or *Stappia* co-cultures in either dust treatment, indicating that they were either produced directly by *Marinobacter* or by *P. tricornutum* only when *Marinobacter* was present. These compounds were more abundant in the low dust treatment, indicating that they may have been produced due to Fe limitation stress. In contrast, the one detected feature that was unique to the *Stappia* co-cultures (C_17_H_36_O_4_S_2_) was most abundant in the high dust treatment. This compound is likely not tied to Fe limitation stress, but rather to *Stappia*’s abundance, as bacterial cell counts were 31 times higher in the high dust *Stappia* co-cultures than in the low dust treatment.

The 5 features associated with bacterial presence (C_9_H_12_O_2_N_4,_ C_25_H_44_O_4_S_2_, C_12_H_18_O_2_N_4_, C_8_H_6_O_2_S, C_15_H_14_O_2_S) and the 6 diatom density-associated features that were more abundant in the presence of bacteria (C_14_H_29_O_3_N, C_13_H_14_O_4_S, C_12_H_18_O_2_N_4_, C_12_H_16_O_2_N_4,_ C_12_H_16_O_3_N_4,_ C_12_H_20_O_5_N_4_) tended to be classified as peptide-like or organosulfur-type compounds, with a higher proportion of organosulfur features belonging to the bacteria-associated compound cluster. Compounds in the bacteria associated cluster tended to be more abundant in the low dust treatment, indicating that these compounds were likely produced in response to Fe limitation stress.

Exometabolites associated with higher algal cell density included purine nucleoside-type and indole-like compounds, including MS/MS annotated deoxyadenosine (C_10_H_13_O_3_N_5_), adenosine (C_10_H_13_O_4_N_5_), indolactic acid (C_11_H_11_O_3_N), indole-3-acetaldehyde (C_10_H_9_ON), and 3-formylindole (C_9_H_7_ON) (Table 1, Fig. S5). This cluster also includes 4 features assigned molecular formula (C_20_H_41_O_4_N, C_15_H_24_O_3_N_4_S_2,_ C_6_H_7_ON_5_, C_2_H_4_O_3_N) that were not structurally annotated.

## Discussion

Despite the low bacterial abundance in the initial experiment with algal cells outnumbering bacterial cells 20:1 after 14 days in co-culture with *Marinobacter* and 16:1 in co-culture with *Stappia*, bacterial presence had a significant effect on *P. tricornutum*’s growth. The results of this study indicate that low abundance members of the bacterial community may have a disproportionate impact on algal growth. Notably, the growth enhancement effect of *Marinobacter* and inhibitory effect of *Stappia* in this study were only observed in the low dust treatment where Fe was the limiting nutrient. This suggests that the bacterial growth effect on *P. tricornutum* were related to increased Fe availability.

The rapid growth response of *P. tricornutum* in co-culture with *Marinobacter* after dust addition indicates that *Marinobacter* is effective at mobilizing Fe from insoluble mineral phases to promote algal growth. Furthermore, the faster initial growth of *Marinobacter* in the co-cultures under low dust conditions compared to high dust conditions suggests that *P. tricornutum*, which is supported by the rapid increase in bacterial cells in the *P. tricornutum-Marinobacter* co-culture grown under low dust conditions. This is consistent with the timescale of natural community responses to dust deposition under variety of nutrient limitations (e.g., N, P, Fe), with several studies reporting an increase in heterotrophic biomass [63,64] and primary production in days after dust deposition events. Likewise, chlorophyll a has been reported to increase within 0-5 days of dust input in incubation [65–68] and remote observation [69] studies alike. While the experiments conducted during this study showcase interactions between a single diatom and bacterial strain, there is some evidence that dust deposition events with natural communities may increase interspecies interactions [68], so developing an understanding of the mechanisms of these interactions is critical for understanding and predicting system-level responses to such events.

The mechanism of bacterial effects on diatom growth and Fe solubilization remain unclear, but genomic evidence for the capacity to produce Fe-chelating siderophores suggests that these pathways may be important. Although *P. tricornutum* cannot produce siderophores, it possesses an uptake receptor FBP1 and ferric reductase FRE2 that are upregulated under Fe deficiency and potentially involved in bacterial siderophore iron acquisition [12,26]. Since siderophores are structurally diverse and uptake receptors are selective for particular structures [70], bacterial siderophores that solubilize iron from dust could potentially have beneficial or inhibitory effects on diatom growth depending on whether they can be utilized.

While no Fe-bound complexes were detected in the co-cultures, both *Marinobacter* and *Stappia* were suspected to produce Fe chelators based on predicted biosynthetic clusters (Fig. S1). In a separate experiment comparing Fe-EDTA deficient and replete co-cultures with *Stappia*, a potential siderophore candidate was detected as an Fe-bound complex (C_23_H_32_O_4_N_2_S_2_Fe^+^, *m/z* 520.115; Fig. S6) by LC-MS. It is possible that this compound was below detection limits given the low bacterial cell density in the low dust *Stappia* co-cultures (1.9×10^3^ cells/mL), but was still produced and facilitated Fe-uptake by *Stappia*. Indeed, siderophores are frequently present at picomolar in marine waters [51,71–73], and these low concentrations are likely enough to sustain microbial growth. If this compound was not compatible with *P. tricornutum* uptake mechanisms, this could explain the observed algal growth suppression. For *Marinobacter*, the 2 metabolites (C_11_H_23_O_9_N and C_17_H_18_O_4_S) associated solely with the low-dust *Marinobacter* co-cultures are targets of interest with a potential role in solubilizing mineral-bound Fe. It is possible that these molecules, or other exometabolites from *Marinobacter*, solubilize Fe in a form that is bioavailable to *P. tricornutum*. Structural categorization of these molecules and their interactions with iron minerals would shed light on their potential role in the partnership between *P. tricornutum* and *Marinobacter* as well as in Fe mobilization.

It is also possible that Fe was mobilized by *Marinobacter* from the mineral dust in this experiment by a reductive dissolution mechanism [74–76]. Though Fe(II) measurements were conducted as part of the dust addition experiments to attempt to capture elevated concentrations after dust addition, all measurements were close to the detection limit of the method (25 pM) [77,78]. However, these results do not preclude a reductive dissolution mechanism since Fe(II) oxidation and uptake kinetics can be rapid [79–82]. Future work characterizing *Marinobacter* and *Stappia*’s potential to colonize and solubilize Fe-bearing minerals will help to clarify the mechanism of Fe-limited algal growth enhancement and inhibition for each respective bacterial strain.

Other abundant metabolites significantly enhanced by *Marinobacter*’s presence in the low dust treatment included indoles structurally related to known auxins that promote *P. tricornutum* growth. Indoles such as indole acetic acid (IAA) serve as plant growth hormones and signaling molecules which can increase algal growth [83–87], and have been observed in *P. tricornutum* exudate in previous studies [38]. Though IAA was not detected in this study, indole-3-acetaldehyde may be converted to IAA through an enzymatic pathway [83,86]. It is possible that *P. tricornutum* growth was stimulated by the presence of the detected indoles or their derivatives under the low Fe growth conditions, although these compounds may have alternatively accumulated because of the increased *P. tricornutum* cell density. Despite the known role of indoles in promoting algal growth, it is unlikely that they were fully responsible for *Marinobacter*’s growth enhancement effect observed in this study, especially since these compounds were abundant in the presence of the bacteria under the high dust treatment yet no significant effect of *Marinobacter* was observed when Fe was not limiting.

Differing quotas and acquisition strategies for micronutrients such as Fe [88–90] are a significant driver of phytoplankton community structure in marine ecosystems. Metanalysis of *Tara* Oceans datasets has suggested that sub-assemblages within marine planktonic communities may respond differently to Fe limitation [88]. While photoautotrophs in marine ecosystems have their own strategies for dealing with micronutrient limitation, such as altering gene expression to reduce cellular Fe requirements [29,91], the role of bacterial or low-abundance partners in alleviating micronutrient stress should not be overlooked. Metagenomic analyses of bacterial communities in Fe-deficient ocean regions have revealed heterotrophic bacterial taxa with the capacity to produce structurally diverse siderophores [19,71]. While these taxa typically constitute a small fraction of the overall marine community, the results of our study suggest that they may have outsized impacts on mineral-bound iron accessibility that could impact overall primary production.

As natural dust-derived Fe delivery to marine systems is expected to change under projected climate change scenarios [92] and the scientific community re-visits artificial Fe fertilization as a geo-engineering strategy to mitigate atmospheric CO_2_ increases [93,94], understanding how marine communities can solubilize mineral-bound forms of Fe is more important than ever. There is also growing interest in manipulating algal microbiomes to improve algal production for industrial and biofuel applications [95,96]. Being able to predict which members of a community control Fe availability may enable microbiome manipulation through controlled added Fe sources. In this study, we have demonstrated that low abundance bacteria could be important in moderating algal Fe limitation, particularly when Fe is present in an environmentally-relevant form such as dust. Understanding community-level responses to micronutrient limitation and the potentially outsized role of low-abundance taxa in moderating that stress is critical for improving our ability to predict algal growth and resilience in both natural and engineered systems.

## Acknowledgements

The authors would like to thank Kathleen Kouba for assistance and support with algal culture maintenance. We would also like to thank Dr. Chih-Ping Lee for assistance and training in flow cytometry and Dr. Stephen Giovannoni for access to lab facilities and the Guava Technologies flow cytometer (Millipore; Billerica, MA), as well as Dr. James Moffett and Mary Kate Dinneen for assistance with Fe(II) measurements.

## Author Contributions (CRediT Roles)

NR Coffey: Writing – review & editing, Writing – original draft, Visualization, Validation, Methodology,

Investigation, Formal analysis, Data curation

BN Newell: Writing - review & editing, Investigation

K Manning: Writing - review & editing, Investigation

KA Rolison: review & editing, Investigation

X Mayali: Writing – review & editing

RK Stuart: Writing – review & editing, Project administration, Funding acquisition, Conceptualization

RM Boiteau: Writing – review & editing, Writing – original draft, Supervision, Software, Methodology, Funding acquisition, Formal analysis, Data curation, Conceptualization

## Supplementary Material

Supplementary information is available at The ISME Journal online.

## Competing Interests

The authors declare no competing interests.

## Funding

This this research was funded by U.S. Department of Energy (DOE) Office of Biological and Environmental Research, SCW1039. The work at LLNL was performed under DOE auspices of the by Lawrence Livermore National Laboratory, United States, under Contract DE-AC5207NA27344.

## Data Availability Statement

MS data is available in the MassIVE repository (accession #MSV000102217).

## Notes

https://doi.org/doi:10.25345/C5TH8C232

### Competing Interest Statement

The authors have declared no competing interest.

