## Supplemental Information for "Phycosphere-associated bacteria differentially impact accessibility of dust-bound iron to model diatom *Phaeodactylum tricornutum*"

**Supplemental Methods - Chemiluminescent Fe(II) Measurement**:

Dissolved Fe(II) measurements were conducted following standard methods [1, 2]. Briefly, a concentrated luminol reagent was prepared by combining 254 mL optima grade ammonium hydroxide with 0.796 g luminol, 705 mL ultrapure water, and 42 mL concentrated HCl (Aristar plus) in an acid-cleaned amber HDPE bottle. A working reagent was created by diluting the concentrated reagent 4x with ultrapure water in an acid-cleaned amber HDPE bottle and baking for 12 hours at 50°C. Fe(II) was measured with a luminol chemiluminescence-based method using an FeLume flow-injection system (Waterville Analytical) and Waterville Analytical software. Samples and reagents were introduced using a peristaltic pump at a flow of 2.1 mL/min, and combined in a standard quartz flow cell. Resultant chemiluminescence was quantified using a Hamamatsu HC135 photon counter with an integration time of 200 ms, with data collection occurring for up to 200 s. All samples were introduced in trace metal-clean polypropylene centrifuge tubes.

**Supplemental Figures**:

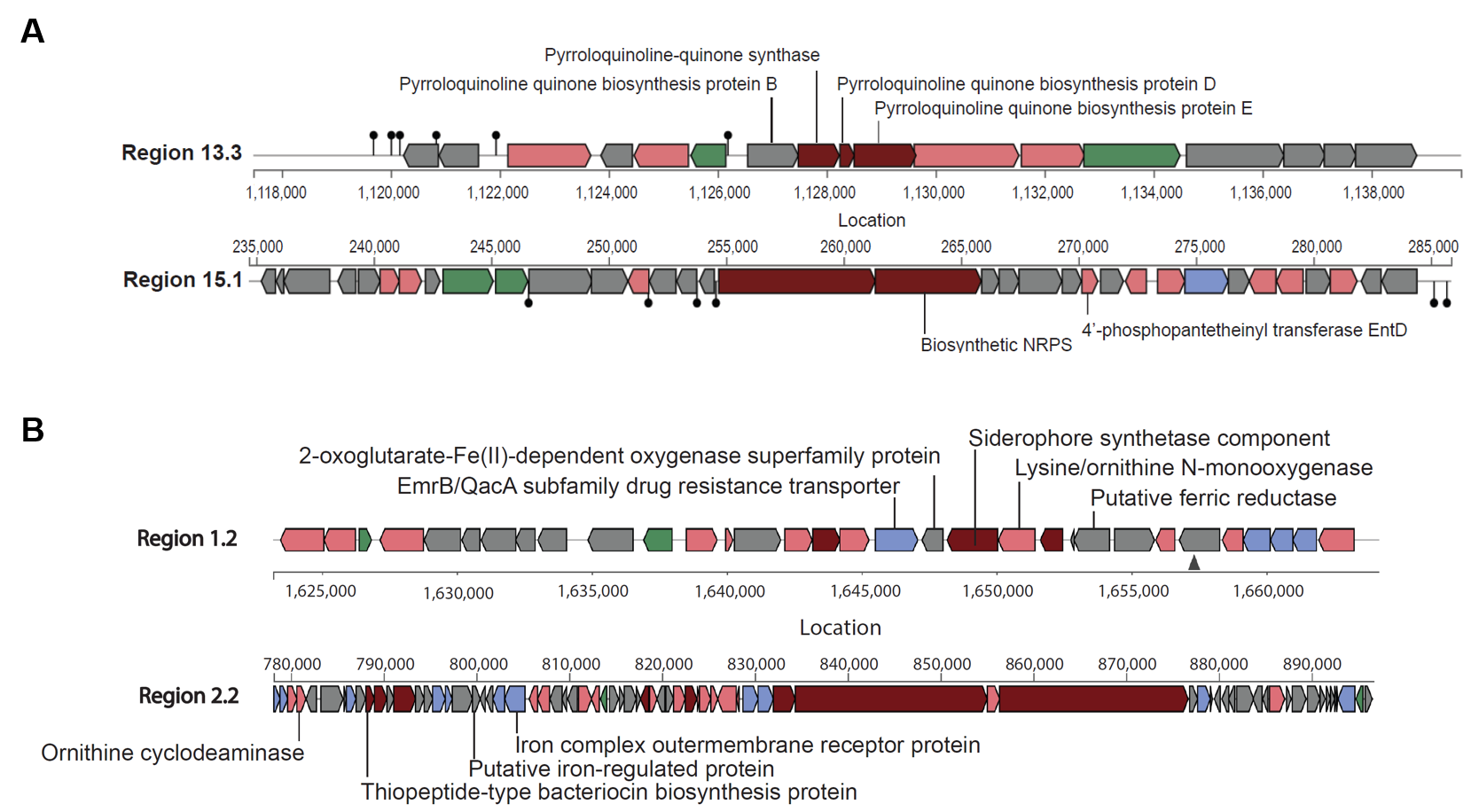

Figure S1: antiSMASH genome annotations of regions of interest for A) *Marinobacter* 3-2, indicating the genomic potential for redox shuttle (Region 13.3) or siderophore (Region 15.1) production; and B) *Stappia sp*. ARW1T, indicating the genomic potential for siderophore biosynthesis.

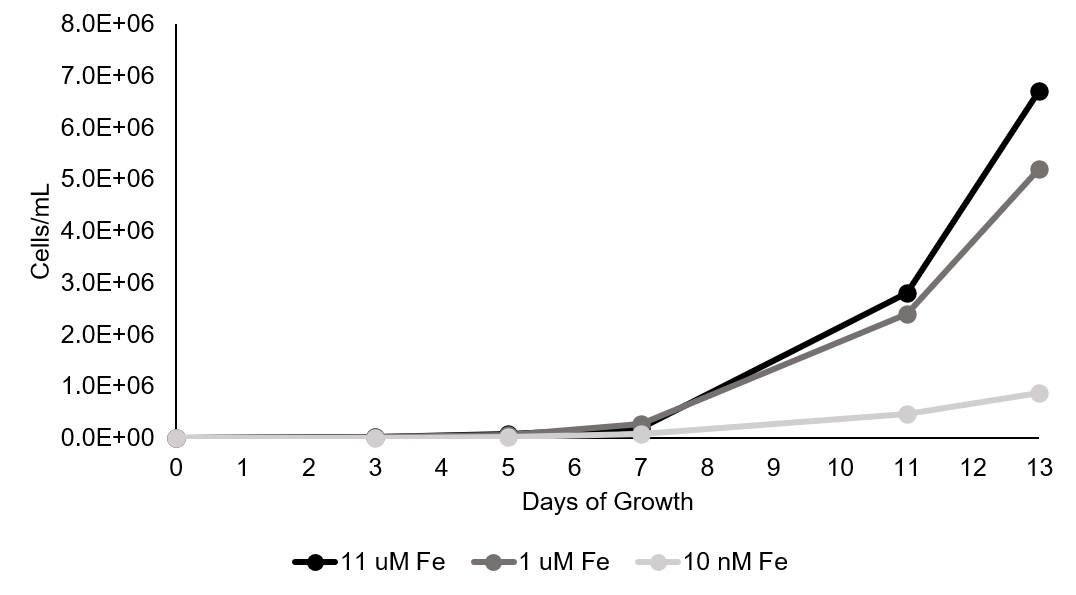

Figure S2: Growth curve of *P. tricornutum* with standard f/2 +Si Fe concentration (11.2 μM), 1 μM Fe, and 10 nM Fe (all conditions with FeCl_3_ added to media containing an excess of EDTA). 10 nM Fe effectively limits *P. tricornutum*’s growth, while 1 μM Fe is more reflective of an Fe-replete condition.

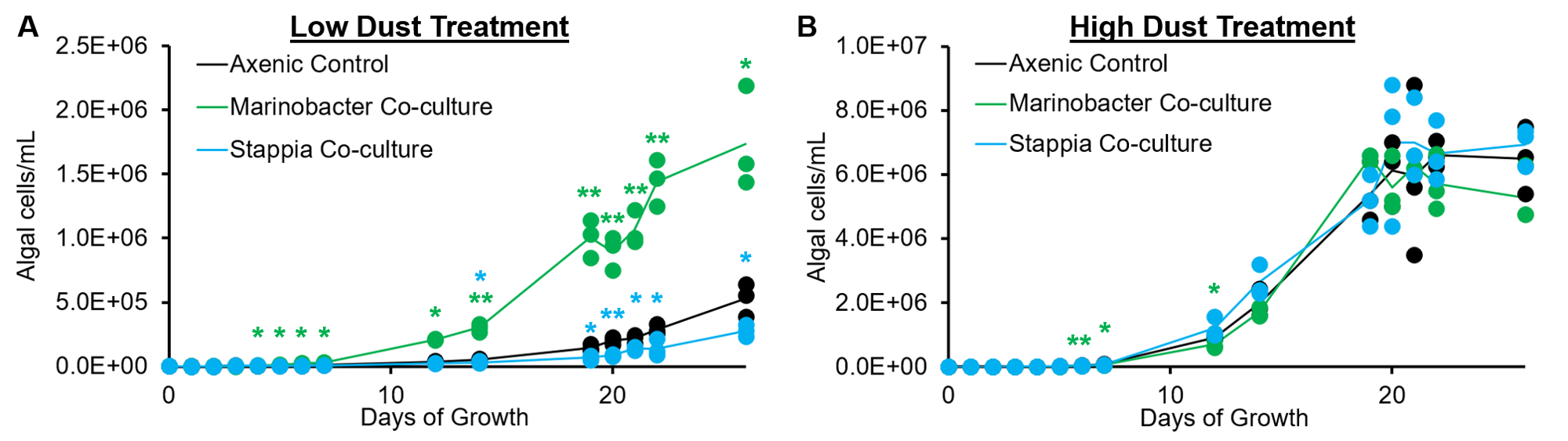

Figure S3: Algal cell density (cells/mL) in low (A) and high (B) dust treatments of axenic *P. tricornutum* (black) and in co-culture with *Marinobacter* (green) and *Stappia* (blue). Individual replicates are plotted as points, while the line represents the average of 3 replicates for each treatment. Significant differences between algal cell abundance with and without bacteria (unpaired heteroscedastic t-test) are noted marked “*” for p < 0.05 and “**” for p < 0.01.

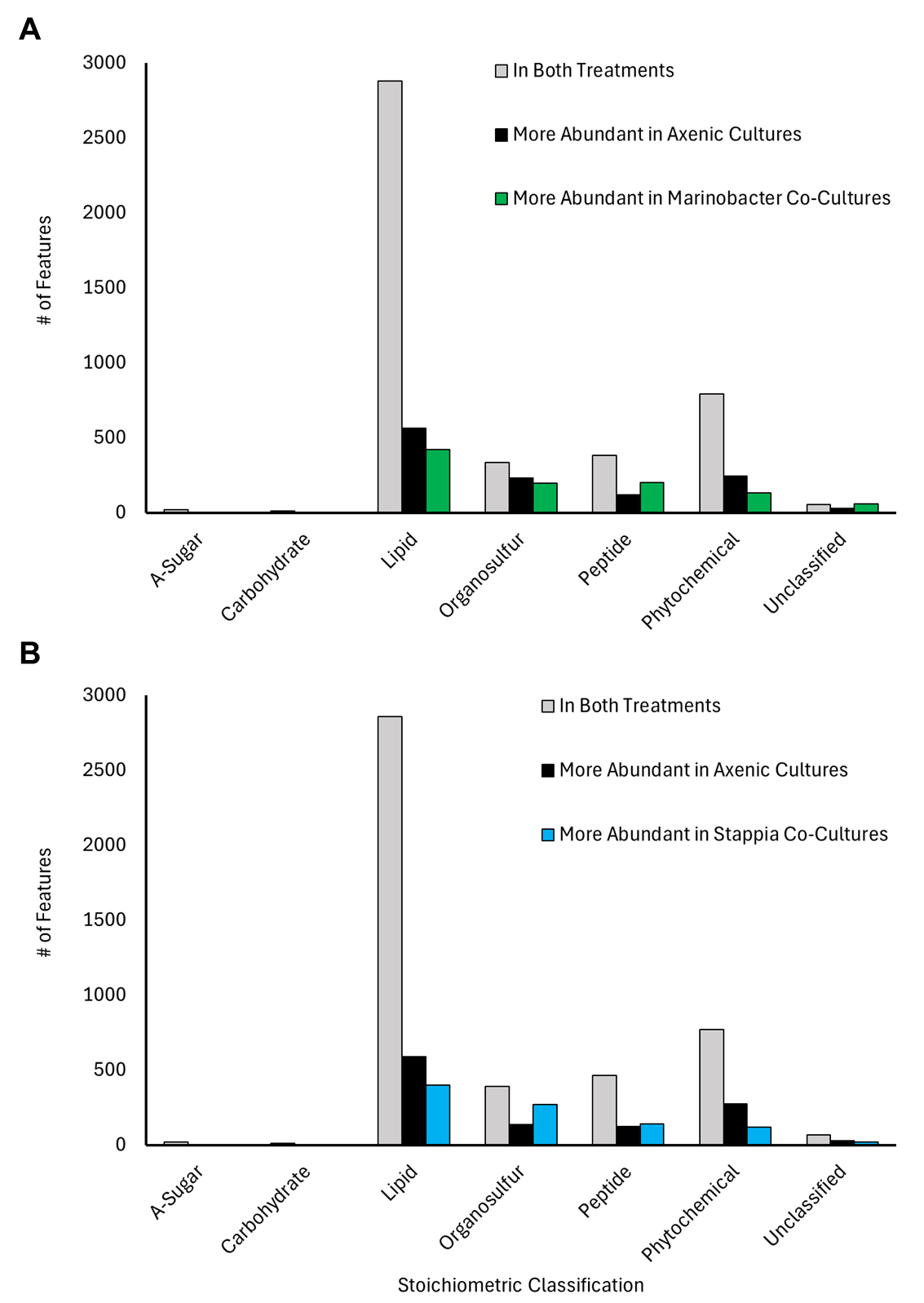

Figure S4: Bar charts showing the number of exometabolome features detected in low dust treatments after 14 days of growth associated with different stoichiometric classifications that were (A) contained in both the axenic diatom culture and in co-culture with *Marinobacter* (grey), ≥2-fold more abundant in the axenic diatom culture (black), or ≥2-fold more abundant in co-cultures with *Marinobacter* (green); or B) contained in both the axenic diatom culture and in co-culture with *Stappia* (grey), ≥2-fold more abundant in the axenic diatom culture (black), or ≥2-fold more abundant in co-cultures with *Stappia* (blue).

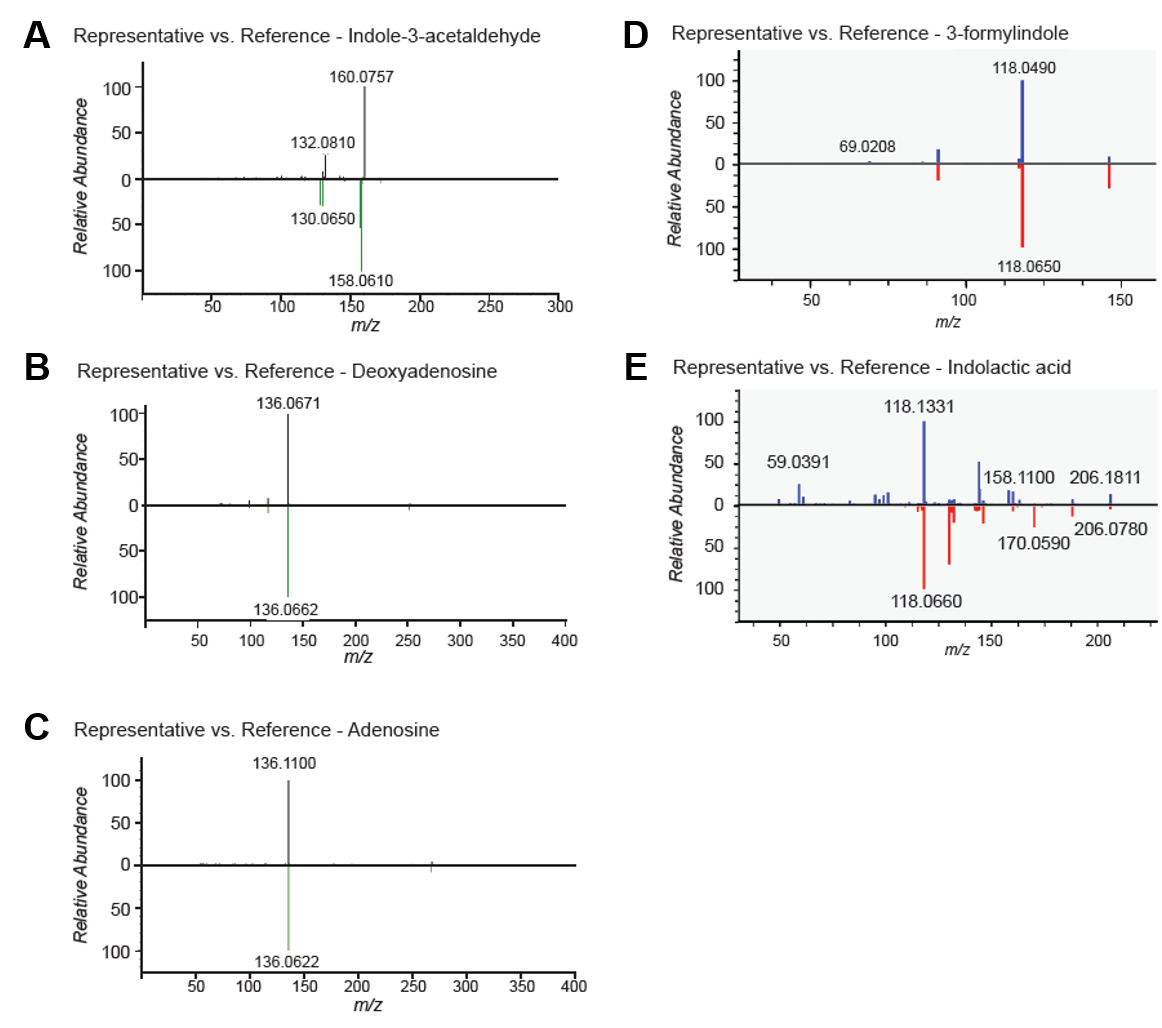

Figure S5: MS/MS spectral mirrors for A) indole-3-acetaldehyde, GNPS2 cosine similarity = 0.8786 (*m/z* tolerance increased to 2.1 due to match to (-) mode library); B) deoxyadenosine, GNPS2 cosine similarity = 0.9990; C) adenosine; GNPS2 cosine similarity = 0.9983; D) 3-formylindole; MSDial match score = 1.8639; and E) indolactoc acid, MSDial match score = 1.212.

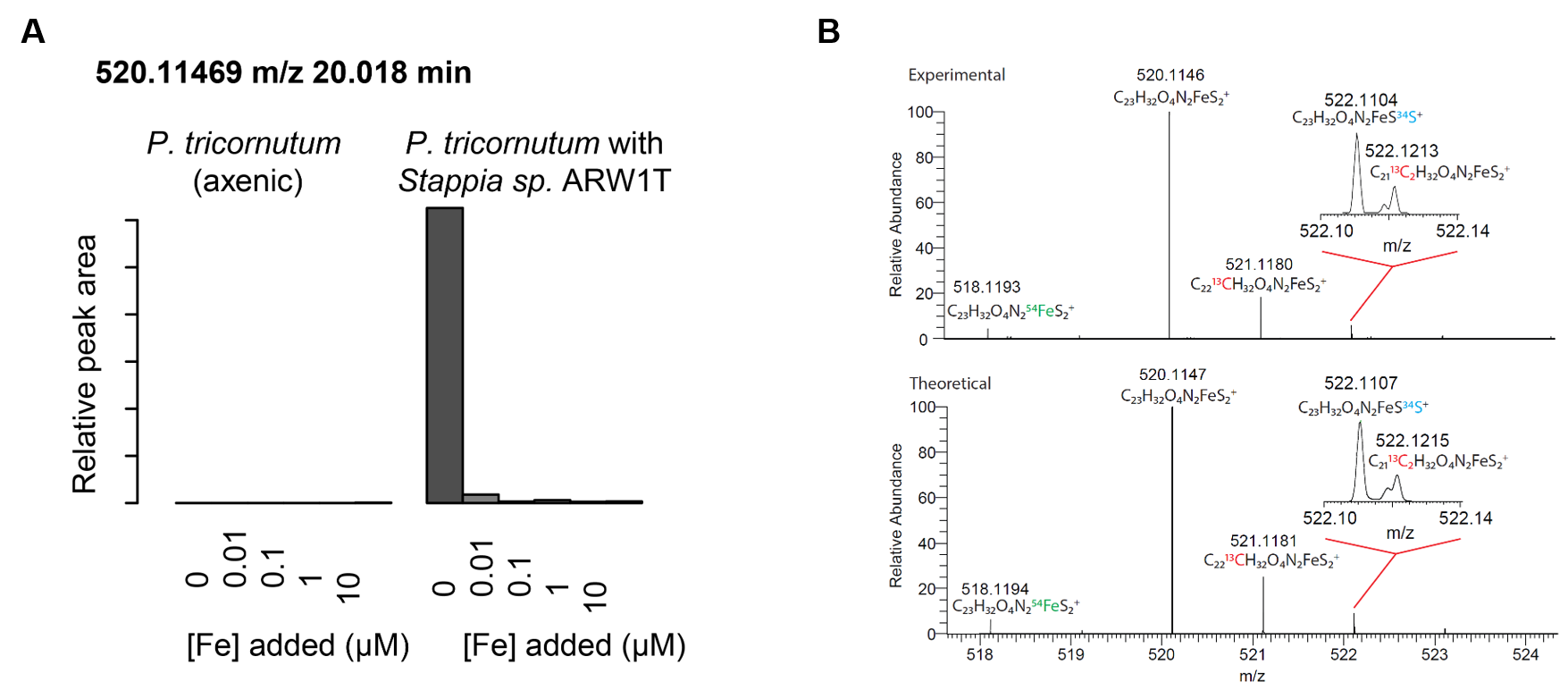

Figure S6: A) Intensity of the putative *Stappia* sp. ARW1T-produced siderophore (*m/z* = 520.1147) across varied Fe-EDTA concentrations in the axenic control and the *P. tricornutum*-*Stappia* co-culture in previous experiments (triplicates pooled for analysis). B) Comparison between the experimentally-determined isotope pattern of the proposed formula (C_23_H_32_O_4_N_2_S_2_Fe^+^) of the putative novel siderophore produced by *Stappia* (top panel) and the theoretical isotope pattern (bottom panel).
